# Evaluating the genetic diagnostic power of exome sequencing: Identifying missing data

**DOI:** 10.1101/068825

**Authors:** Pascual Lorente-Arencibia, Deyán Guacarán, Antonio Tugores

## Abstract

A hurdle of exome sequencing is its limited capacity to represent the entire exome. To ascertain the diagnostic power of this approach we determined the extent of coverage per individual sample. Using alignment data (BAM files) from 15 exome samples, sequences of any length that were below a determined sequencing depth coverage (DP) were detected and annotated with the Ensembl exon database using MIST, a novel software tool. Samples sequenced at 50X mean coverage had, on average, up to 50% of the Ensembl annotated exons with at least one nucleotide (L=1) with a DP<20, improving to 35% at 100X mean coverage. In addition, almost 15% of annotated exons were never sequenced (L=50, DP<1) at 50x mean coverage, reaching down to 5% at 100x. The diagnostic utility of this approach was tested for hypertrophic cardiomyopathy, a genetically heterogeneous disease, where exome sequencing covered as much as 80% of all candidate genes exons at DP≥20. This report stresses the value of identifying, precisely, which sequences are below a specific depth in an individual’s exome, and provides a useful tool to assess the potential and pitfalls of exome sequencing in a diagnostic or gene discovery setting.

**COMPETING INTERESTS:** The authors declare no conflicts of interest.

## INTRODUCTION

Genetic screening for specific variants may predict the susceptibility to suffer disease or the capacity to respond adequately to specific medications, providing a framework to make informed decisions about a clinical strategy. Exome sequencing has allowed to search for disease-causing genetic variants in a hypothesis-free fashion on the most understood fraction of the genome (1-2%), where the biological meaning of a variant might be better inferred ^1^. This strategy has been able to identify disease-associated variants in, approximately, 25% of the probands ^2-4^.

Both conceptual and technical reasons can be argued to explain this incomplete success rate. First, exome sequencing is a gene-centered approach, so regulatory regions are missed ^5^. Secondly, it fails to identify genomic rearrangements, as well as large insertions and deletions (indel). Additionally, even in the best-case scenario, when a variant is a single nucleotide polymorphism or variant or a small indel in an exon, there is still room for failure as (i) not all the exons are represented in the capture kits and (ii) there are sequences that are not sequenced at sufficient depth or at all ^6-9^.

While the overall performance of this analysis has been evaluated at large ^6-10^, a diagnostic approach requires its evaluation on an individual basis, identifying not only the variants, but also the number of protein coding exons that have been missed or sequenced below acceptable quality.

## METHODS

### Subjects and sample sequencing

Fifteen individuals gave written informed consent to participate in three studies of uncharacterized Mendelian disorders by using whole exome sequencing, approved by the Clinical Research Ethics Committee of our Center. DNA was purified from venous blood (QIAGEN), exon-captured with the SureSelect 50Mb kit (Agilent Technologies), and sequenced with a HiSeq2000 (Illumina) as described ^6^. Five samples were sequenced at an average depth of 50 fold (50X) coverage, and the remaining ten samples were re-sequenced to attain a 100 fold (100X) coverage.

### Bioinformatic analysis

Sequence FASTQ files (length = 90 nucleotides) were processed following the Genome Analysis Tool Kit (GATK) best-practices^11^, and aligned to the GRCh37.p13 human reference genome using Burrows-Wheeler Aligner ^12^ with default parameters for paired-end sequences to produce raw SAM files. Sequences which partially fall out of the reference genome were soft-clipped, mapping quality values (MAPQ) were set to 0 in unmapped sequences, duplicated PCR fragments were removed, alignments were sorted for each resulting BAM file using Picard Tools ^13^, indel realignment was performed and recalibrated using GATK ^11^.

MIST (<u>MI</u>ssing <u>S</u>equences detection <u>T</u>ool) is a Java-based software that uses two parameters: **threshold**, which determines the depth (DP) or number of reads to consider for any nucleotide, and **length** (L), which determines the minimum number of consecutive nucleotides under a given DP. The application defines the intervals that have a DP below the given threshold and intersects them with the Ensembl exon database ^15^ including sequences padding the exons (±10 nucleotides) and generates an output file where every line contains a match between an Ensembl exon and a poorly sequenced region (Table 1). MIST source code may be downloaded at: https://github.com/pasculorente/Mist.

**Table 1.**
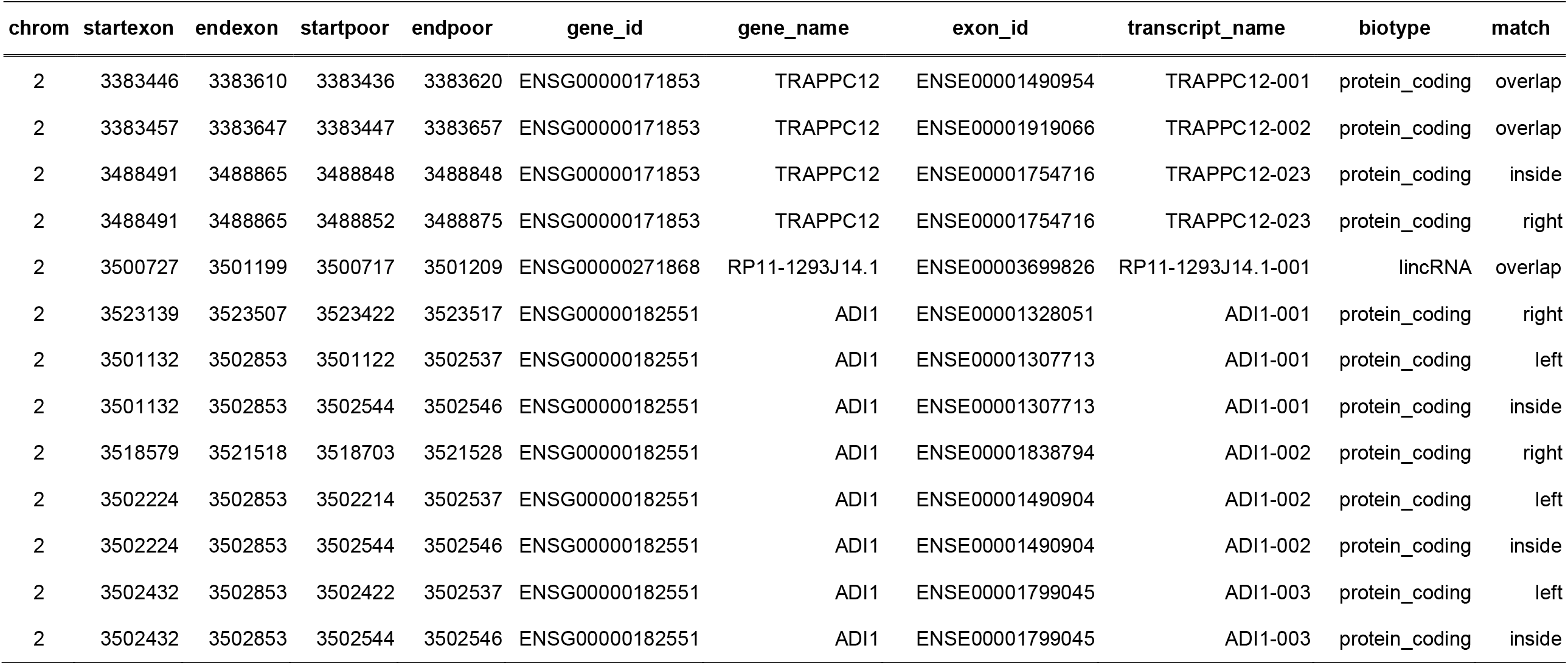
Example of MIST output. Results shown are from a limited number of lines from a single individual. Every region below a user– defined DP is annotated with the following fields: **chrom** (chromosome), **startexon** (the start position of the exon), **endexon** (the end position of the exon), **startpoor** (the start position of the poor region), **endpoor** (the end position of the poor region), **gene_id** (the Ensembl gene identifier), **gene_name** (the Ensembl gene name), **exon_id** (the Ensembl exon identifier), **transcript_name** (the Ensembl transcript name) **, biotype** (the biotype of the exon), **match** (whether the poor region is left, right, inside or overlapping the exon).

## RESULTS

### Sequence coverage of exome sequencing

Five samples were sequenced at an average depth of 50X, and ten at 100X, obtaining 57 to 64 million reads per sample-lane (50X). Sequences present in autosomal or sex chromosomes, and annotated as protein coding exons, containing at least one nucleotide (L=1) below DP=20, were referred as “incomplete” and were identified by their unique Ensembl ID. Sequencing at 50X rendered, on average, over 192,000 incomplete protein-coding exons, roughly representing 50% of the Ensembl-annotated sequences, improving significantly at 100X, where an average of 35% of the Ensembl protein-coding exons were below the threshold (Supplementary Table 1; Figure 1A).

**Figure 1.**
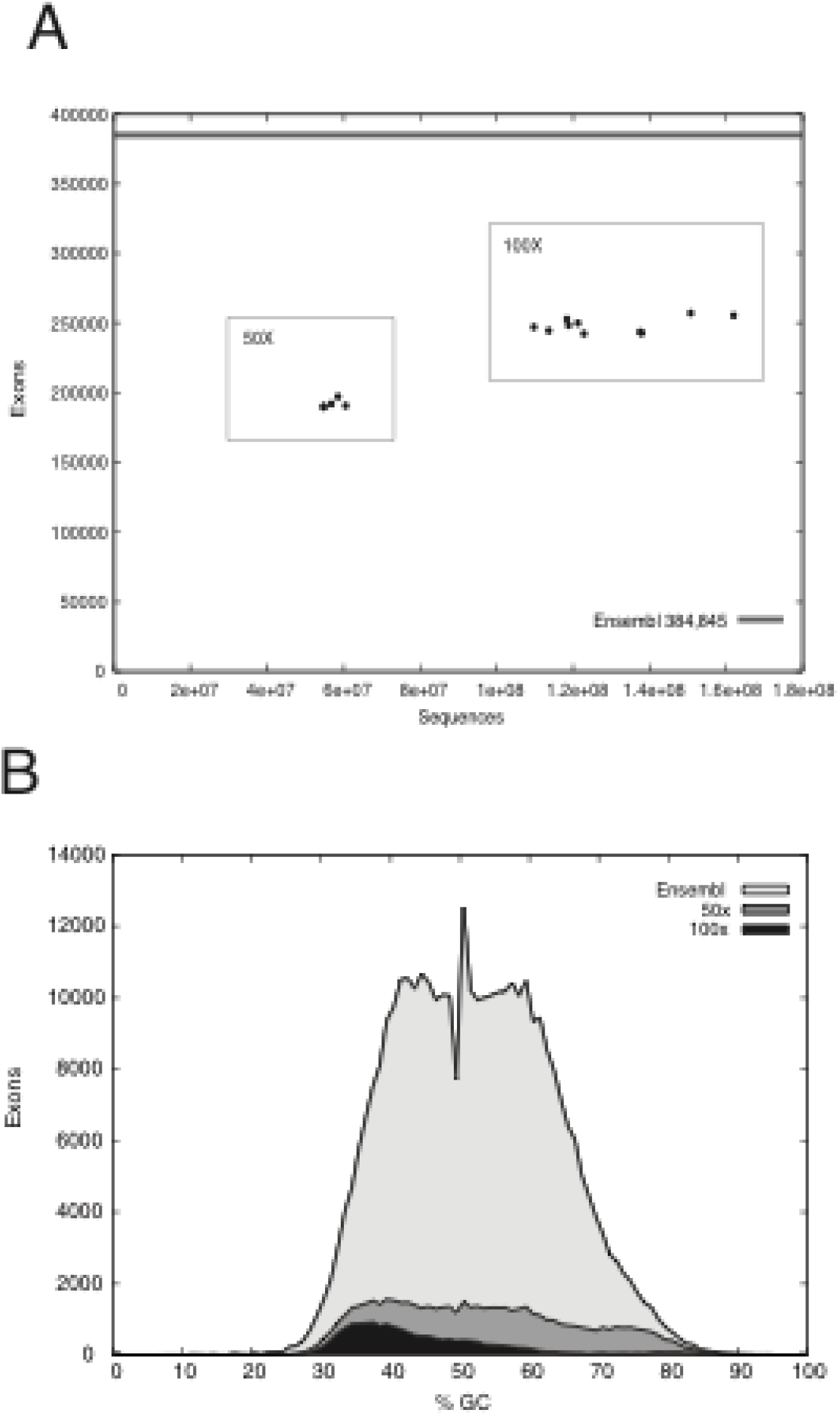
Evaluation of missing sequences in exome sequencing projects. **(A) Representation of the number of reads versus the number of correctly sequenced protein coding exons.**The Y– axis represents the number of complete exons while X–axis represents the number of reads. The 50X and 100X clusters are boxed. The top grey line represents the total number of unique protein coding exons annotated by Ensembl. **(B) “Lost” sequences and G+C content per coverage group.** Protein coding exons that had, at least, a 50 nucleotide stretch that was never read (DP<1), were considered as completely lost or missed by the analysis. The graphic shows the total number of “lost” protein coding exons, as determined by the intersection of of all samples within the 50x (dark grey shading) and 100x (black shading) groups, in comparison with the total number of unique protein– coding exons annotated by Ensembl (light grey shading). The GC content is shown, as percentage, on the Y axis.

The number of sequences that are not represented in the exome capture kits varies depending on the kit, and on the databases used as reference ^6-9^. These sequences should appear as missing in each sample, along with those that are sequenced with poor quality or not at all. A threshold of DP=1 and L= 50 or higher, and the intersection for each group (50X and 100X) identified exons that were consistently missed among all samples from a given group. Almost 15% of the protein-coding exons (56656/384845) were never sequenced in the 50x coverage group while only 5% of the protein coding exons (17838/384845) were absent from the 100X group, showing a lower GC content than average (Figure 1B).

### Exome sequencing in genetic diagnosis

The diagnostic limits of exome sequencing were tested for hypertrophic cardiomyopathy (CMH; MIM: 192600), characterized by great genetic heterogeneity: 22 candidate genes, representing 999 exons present in 96 transcripts ^16^. Out of the list of known candidate genes, exome sequencing was able to detect most exons, revealing that there were, on average, only 169 (17%) and 121 exons (12%) with at least one nucleotide below DP=10, and 254 (25%) and 171 exons (17%) below DP=20 (Table 2) in the 50x and 100x sample groups respectively.

**Table 2.** Missing sequences per sample in genes known to be associated with hypertrophic cardiomyopathy. MIST software was run on each individual sample to detect sequences of at least 1 nucleotide in length with thresholds of DP<10 and DP<20. Data was filtered, for each individual sample, to include only unique exons annotated in Ensembl as part of genes that have been associated with hypertrophic cardiomyopathy according to the OMIM database at the National Center for Biotechnology Information (NCBI; http://www.ncbi.nlm.nih.gov/). Numbers shown for individual samples define exons that have at least one nucleotide with a DP<20.

| Disease | MIM | Gene | Ex.N | N019 |  | N032 |  | N060 |  | W002 |  | S001 |  | S020 |  |
| --- | --- | --- | --- | --- | --- | --- | --- | --- | --- | --- | --- | --- | --- | --- | --- |
|  |  |  |  | 10 | 20 | 10 | 20 | 10 | 20 | 10 | 20 | 10 | 20 | 10 | 20 |
| CMH1 | 192600 | CAV3 | 8 | 3 | 4 | 2 | 4 | 3 | 3 | 2 | 2 | 2 | 2 | 2 | 2 |
| CMH1 | 192600 | MYH7 | 40 | 3 | 9 | 4 | 14 | 4 | 10 | 6 | 12 | 6 | 12 | 5 | 10 |
| CMH1 | 192600 | MYLK2 | 20 | 5 | 10 | 6 | 9 | 5 | 7 | 2 | 3 | 2 | 3 | 1 | 2 |
| CMH2 | 115195 | TNNT2 | 81 | 16 | 19 | 15 | 17 | 13 | 17 | 5 | 8 | 5 | 14 | 4 | 8 |
| CMH3 | 115196 | TPM1 | 85 | 28 | 32 | 29 | 34 | 28 | 28 | 12 | 16 | 10 | 11 | 9 | 10 |
| CMH4 | 115197 | MYBPC3 | 39 | 16 | 23 | 19 | 26 | 14 | 23 | 3 | 6 | 3 | 6 | 3 | 3 |
| CMH6 | 600858 | PRKAG2 | 64 | 15 | 17 | 14 | 17 | 15 | 16 | 9 | 9 | 9 | 9 | 9 | 10 |
| CMH7 | 613690 | TNNI3 | 31 | 8 | 11 | 8 | 11 | 8 | 11 | 1 | 4 | 2 | 3 | 0 | 3 |
| CMH8 | 608751 | MYL3 | 13 | 5 | 6 | 6 | 7 | 6 | 10 | 4 | 5 | 4 | 5 | 4 | 5 |
| CMH9 | 188840 | TTN | 387 | 17 | 46 | 13 | 38 | 18 | 45 | 33 | 53 | 33 | 48 | 33 | 42 |
| CMH10 | 160781 | MYL2 | 14 | 2 | 3 | 2 | 4 | 3 | 3 | 1 | 1 | 1 | 2 | 1 | 1 |
| CMH11 | 612098 | ACTC1 | 13 | 2 | 3 | 2 | 3 | 2 | 3 | 2 | 5 | 2 | 4 | 2 | 4 |
| CMH12 | 612124 | CSRP3 | 8 | 4 | 6 | 4 | 5 | 4 | 5 | 4 | 4 | 4 | 4 | 4 | 4 |
| CMH13 | 613243 | TNNC1 | 11 | 3 | 5 | 3 | 6 | 3 | 5 | 1 | 3 | 2 | 2 | 1 | 2 |
| CMH14 | 613251 | MYH6 | 54 | 4 | 11 | 5 | 13 | 4 | 9 | 7 | 12 | 8 | 12 | 6 | 10 |
| CMH15 | 613255 | VCL | 37 | 8 | 12 | 7 | 14 | 9 | 14 | 5 | 9 | 4 | 8 | 5 | 7 |
| CMH16 | 613838 | MYOZ2 | 6 | 2 | 3 | 2 | 2 | 2 | 2 | 2 | 2 | 2 | 2 | 2 | 2 |
| CMH17 | 613873 | JPH2 | 8 | 7 | 7 | 7 | 7 | 7 | 7 | 6 | 7 | 4 | 6 | 5 | 6 |
| CMH18 | 613874 | PLN | 2 | 2 | 2 | 2 | 2 | 2 | 2 | 2 | 2 | 2 | 2 | 2 | 2 |
| CMH19 | 613875 | CALR3 | 14 | 2 | 3 | 3 | 3 | 2 | 3 | 1 | 1 | 0 | 1 | 0 | 1 |
| CMH20 | 613876 | NEXN | 36 | 9 | 10 | 7 | 11 | 7 | 11 | 9 | 9 | 9 | 9 | 9 | 9 |
| CMH22 | 615248 | MYPN | 28 | 8 | 12 | 9 | 12 | 9 | 15 | 8 | 11 | 8 | 11 | 8 | 10 |
| Ensembl total |  | 22 | 999 | 169 | 254 | 169 | 259 | 168 | 249 | 125 | 184 | 122 | 176 | 115 | 153 |

Furthermore, analysis of the protein-coding exons contained the 27 Megabase region linked to CMH21 (MIM 614676; 115 genes, 1864 exons) ^17^, the analysis revealed that 57% and 75% of all annotated exons were fully sequenced at a DP≥10 in N019 (50X) and S003 (100X) respectively,decreased accordingly to 43% and 54% at DP≥20.

## DISCUSSION

In a diagnostic sequencing analysis it is important to know the extent of representation of the genes of interest for a particular disease, regardless of the reason why these genes are absent. Exome capture and sequencing are both physico-chemical processes subjected to varied efficiency, so knowing the un-captured moiety is insufficient to assess missing sequences or below a necessary quality. We found that, as the number of reads increased from 50X to 100X, the number of incomplete sequences decreased. However, a higher number of reads in the 100x sample group did not result in an significantly higher number of fully sequenced exons above the threshold, suggesting that the system could be close to saturation. Moreover, the numbers of commonly shared incomplete or missed protein coding exons were quite close: 118,943 at 50X vs 114,287 at 100X, indicating that missing sequences due to lack of capture and sequencing errors were not significantly rescued by increasing the average coverage at this depth even though it increased the overall quality of the reads. It remains to be determined whether higher average coverage using more sequencing lanes would result in increased detection. We also observed that missing sequences were not due to amplification bias against sequences with high G+C content ^18^.

Exome sequencing has slowly moved into the clinical diagnostic laboratory: Diseases with great genetic heterogeneity with many candidate genes or when not all the possible candidate genes have been identified may benefit from this approach, and has been recently recommended to be part of an integral genetic diagnostic workflow ^3,4,19^. Nonetheless, in addition to the challenges of interpretation of the variants that are found, it is also important to identify the relevant sequences that are incomplete or missing for a specific patient, so that the limitations of the test can be fully understood. In this scenario, we ascertained the degree of coverage for a group of genes associated with CMH (MIM: 192600), a genetically heterogeneous disease, finding that exome sequencing was able to detect up to 83% and 88% of the exons at DP>10 and, and as much as 75% and 83% at DP>20 at 50x and 100x average coverage respectively. This example not only highlights the significant improvement resulting from increasing the average coverage from 50x to 100x but, also, identifies exactly the regions where directed Sanger sequencing is needed.

## ACKNOWLEDGEMENTS

We thank the patients for their participation. Drs. Sarah Murray and Gary Hardiman provided useful comments on the manuscript. Dr Francisca Quintana’s contribution has been pivotal to create and enable the student exchange program between the Hospital and the School of Informatics and Computer Engineering from the University of Las Palmas de Gran Canaria. Work supported by the Fundación Canaria de Investigación en Salud (FUNCIS), and by the Servicio Canario de Salud.

---

**Supplementary Table 1.**
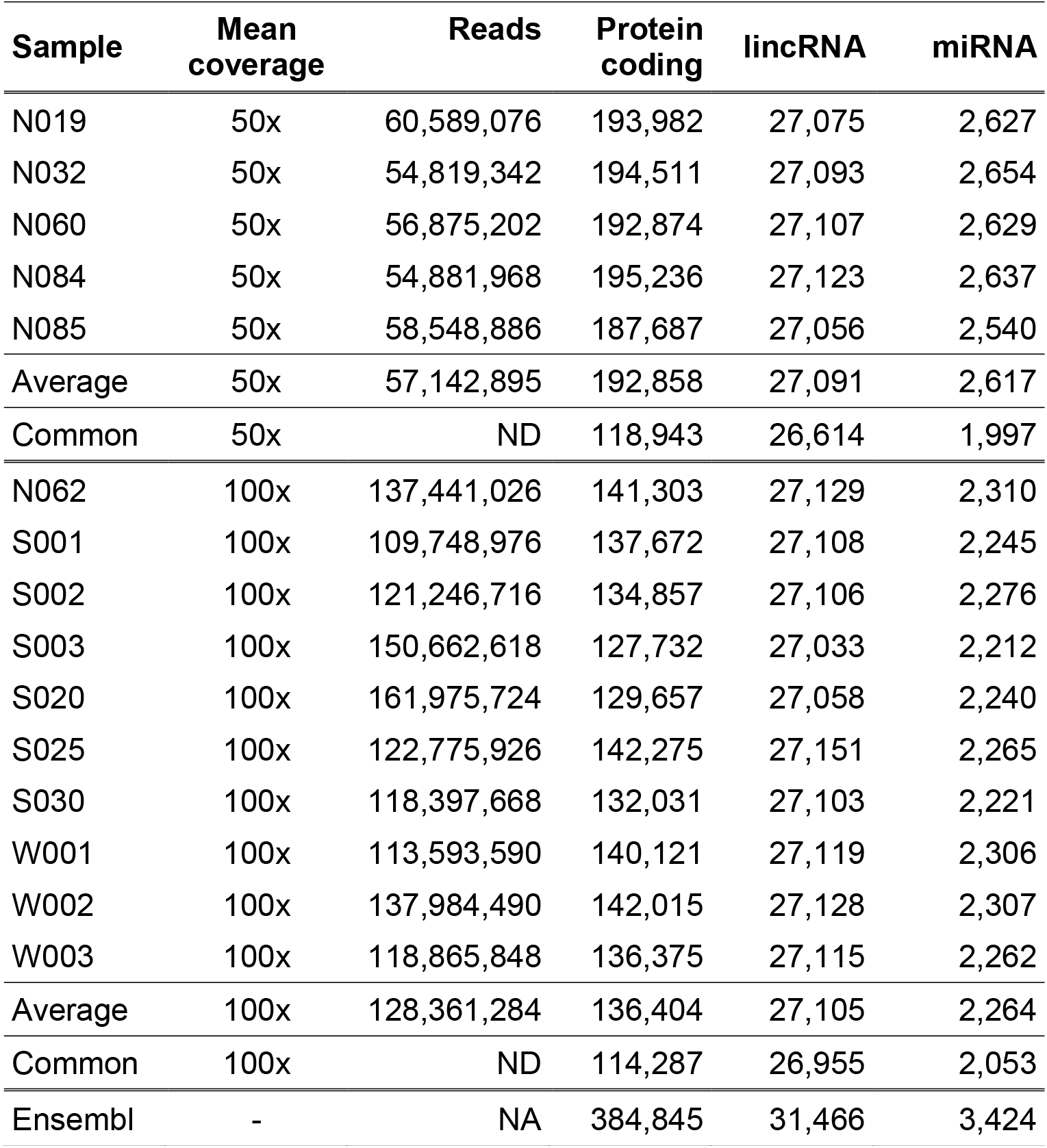
Sequence reads per sample. Genomic DNA isolated from 15 individuals were subjected to exon selection with the Agilent SureSelect 50Mb capture kit and sequencing with an Illumina HiSeq2000. Samples were resolved in one (50x average coverage), or 2 lanes (100x average coverage) as indicated. Final number of sequences is shown for each sample, and the average number per group. after removal of low quality sequences and PCR duplicates, as described in the Methods section. MIST software was run on each individual sample with a threshold of DP≤20 read and a length 1 nucleotide or more. Data was filtered, for each individual sample, to include only sequences present in autosomes or sex chromosomes, excluding mitochondrial and other sequences not assigned to specific chromosomes, according to the annotation of protein coding exons, lincRNAs, and miRNAs in Ensembl. Only unique exons are counted, according to their Ensembl exon ID.

